# Transcriptional Regulators of the *Golli/Myelin Basic Protein* Locus Integrate Additive and Stealth Activities

**DOI:** 10.1101/2020.04.03.023473

**Authors:** Hooman Bagheri, Hana Friedman, Kathy Siminovitch, Alan Peterson

## Abstract

Myelin is composed of plasma membrane spirally wrapped around axons and compacted into dense sheaths by myelin associated proteins. In the central nervous system (CNS), myelin is elaborated by neuroepithelial derived oligodendrocytes and in the peripheral nervous system (PNS) by neural crest derived Schwann cells. While some myelin proteins are unique to only one lineage, *myelin basic protein* (*Mbp*) is expressed in both. Overlapping the *Mbp* gene is *Golli*, a transcriptional unit that is expressed widely both within and beyond the nervous system. A super-enhancer domain within the *Golli/Mbp* locus contains multiple enhancers shown previously to drive reporter construct expression specifically in oligodendrocytes or Schwann cells. In order to determine the contribution of each enhancer to the *Golli/Mbp* expression program and examine if interactions among these enhancers occur, we derived mouse lines in which enhancers were deleted, either singly or in different combinations, and relative mRNA accumulation was measured at key stages of development.
Although super-enhancers have been shown to facilitate interaction among their component enhancers, the enhancers investigated here demonstrated functions that were largely additive. However, enhancers demonstrating autonomous activity strictly in one cell lineage, when missing, were found to significantly reduce output in the other thus revealing cryptic “stealth” activity. Further, *Golli* accumulation in all cell types investigated was markedly and uniformly attenuated by the absence of a key oligodendrocyte enhancer. Our observations expose a novel level of enhancer interaction and are consistent with a model in which enhancer-mediated DNA looping underlies higher-order *Golli/Mbp* regulatory organization.

**AUTHOR SUMMARY:** The control of transcription is mediated through regulatory sequences that engage in a lineage and developmentally contextual manner. The *Golli/Mbp* locus gives rise to several mRNAs and while *Mbp* mRNAs accumulate exclusively in the two glial cell types that elaborate myelin, *Golli* mRNAs accumulate in diverse cell types both within and beyond the nervous system. To determine how the different *Golli/Mbp* enhancers distribute their activities and to reveal if they operate as autonomous agents or have functionally significant interactions with each other we derived multiple enhancer knock-out lines. Comparing the developmental accumulation of *Mbp* and *Golli* mRNAs revealed that the autonomous targeting capacity of multiple enhancers accurately predicted their in-situ contributions. Also, they acted in a largely additive manner indicating significant individual autonomy that can be accounted for by a simple chromatin looping model. Unexpectedly, we also uncovered cryptic “stealth” activity emanating from these same enhancers in lineages where they show no autonomous targeting capacity thus providing new insight into the control of lineage specific gene expression.

## INTRODUCTION

Myelin facilitates rapid and energy efficient conduction of electrical signals and when disrupted, as in demyelinating diseases such as Multiple Sclerosis, nervous system function can be severely compromised. In the mouse, myelination initiates between birth and weaning in both the CNS and PNS with peak accumulation of the mRNAs encoding myelin specific proteins reached during the third postnatal week (1, 2). Adaptive changes in myelin volume are also thought to influence circuit properties in the mature nervous system (3, 4). Consequently, the mechanisms controlling myelin synthesis, including regulation of the genes encoding myelin related proteins, are the focus of intense investigation (5).

Myelin basic protein (MBP), a major myelin component, is an intrinsically disordered protein susceptible to multiple post-translational modifications and while largely concentrated in the myelin sheath, it is implicated in a wide array of cellular functions (6). Mice lacking MBP demonstrate a shivering phenotype and while only limited ultrastructural anomalies and a modest decrease in myelin sheath thickness is observed in their PNS (7–10), compact myelin is absent in their CNS (11–13) Notably, a positive correlation between the accumulated level of *Mbp* mRNA and CNS myelin sheath thickness was reported demonstrating that MBP is not only essential but a limiting factor in CNS myelin production (14, 15). The *Golli/Mbp* locus also expresses *Golli* transcripts that initiate far upstream and incorporate various *Mbp* exons. Golli accumulates in diverse lineages both within and beyond the nervous system on cell-type specific developmental expression programs where it has been shown to modulate Ca^2+^ transients (16–22).

Many of the TFs specifying myelinating cell lineages and/or controlling lineage maturation have been identified (5, 23–34). Further, chromatin remodeling and epigenetic changes associated with myelin gene expression have identified numerous features of the chromatin landscape in which such regulatory components operate (33–40).

Motivated by the critical role myelin plays in nervous system function, and the essential and limiting role of MBP in CNS myelin formation, we and others have characterized features of the mechanism controlling transcription of the *Golli/Mbp* locus (1, 2, 26, 33, 34, 41–46). Because *Mbp* is expressed by the two different cell types that elaborate myelin, accumulates in a well characterized post-natal developmental program and has an intimate and unusual association with the widely expressed overlapping *Golli* transcriptional unit, the *Golli/Mbp* locus represents a particularly rich target within which the higher order organization of transcriptional regulators can be revealed. Moreover, a recently defined super-enhancer domain encompasses multiple *Golli/Mbp* enhancers providing a further opportunity to investigate the function of such domains.

Previously, the lineage specificities and developmental programs conferred by *Golli/Mbp*-associated enhancers were assigned using reporter constructs (2, 41, 43, 44, 46–48). However, in such preparations, enhancers are isolated from their normal chromatin environment and often ligated adjacent to each other or directly to a promoter creating novel spatial relationships that may impose or lead to loss of higher-order interactions (49). Therefore, we sought to characterize enhancer contributions in the fully integrated context of the endogenous *Golli/Mbp* locus. Using CRISPR based gene editing we derived lines of mice bearing alleles deleted of one or more enhancers and these were assessed for *Mbp* and *Golli* mRNA accumulation (relative to *Gapdh* mRNA) at key stages of post-natal development. Such KO alleles caused the greatest reduction in mRNA accumulation in the lineage where the deleted enhancer/s demonstrated autonomous targeting activity in reporter constructs. Unexpectedly, some also led to reduced activity in the lineage where they demonstrated no autonomous targeting capacity and we refer to such additional cryptic functions as “stealth” activity. Our observations are consistent with a model in which transcription factor mediated interactions give rise to chromatin looping that contributes to both *Mbp* and *Golli* programing.

## RESULTS

### Enhancer KO lines

Five modules of high interspecies conservation are located 5’ of the *Mbp* start site within a putative super-enhancer domain (Fig. 1A). In previous studies, M1 (which encompasses the M1E enhancer and the contiguous *Mbp* proximal promoter), M3 and M5 were each shown to drive expression in oligodendrocytes while expression in Schwann cells was driven by M4. M2 demonstrated no autonomous activity in either lineage (2, 41–43, 46). In the present investigation, we derived mouse lines homozygous for alleles deleted individually of M3, M4 or M5 or combined KOs of M3/M5 and M1E/M3/M5. Additionally, to explore the potential function of an enhancer subdomain, an allele was derived bearing a partial deletion of M3, M3(225)KO, that was shown previously to upregulate activity in both oligodendrocytes and Schwann cells (42, 43) (Fig. 1B). At key stages of myelination, accumulation of *Mbp* mRNA was analyzed in spinal cord and sciatic nerves while accumulation of *Golli* mRNA was assessed in spinal cord, thymus and retina at multiple ages.

**Figure 1.**
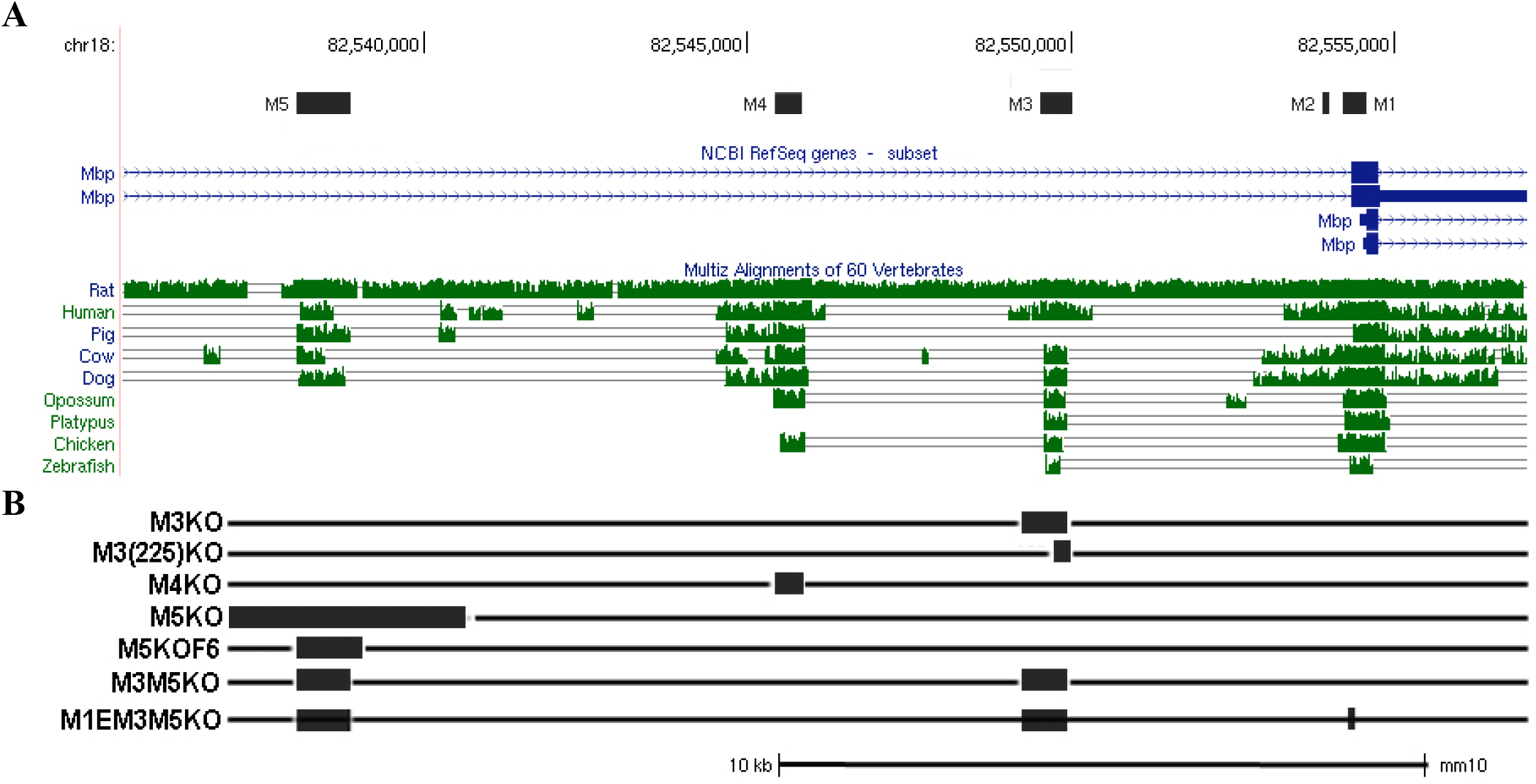
(**A**) *Mbp* 5’ flanking sequence indicating five modules with high interspecies sequence conservation in selected vertebrates (adapted from UCSC browser; see full species comparisons in mouse genome assembly GRCm38/mm10). (**B**) The position and extent of individual and combined enhancer deletions that were derived into individual mouse lines.

### Mbp mRNA accumulation in spinal cord oligodendrocytes

Relative accumulation of *Mbp* mRNA in maturing oligodendrocytes and Schwann cells was measured by analyzing whole tissue homogenates of spinal cord (CNS) and sciatic nerve (PNS) respectively. Oligodendrocytes initiate expression of *Mbp* as a terminal maturation event coincident with myelin sheath elaboration. Although myelination initiates on different schedules in different CNS domains, we restricted our analysis to the cervical spinal cord where myelination initiates perinatally and oligodendrocyte numbers remain constant from at least P10 through P30 (50). Thus, a close relationship between the mRNA levels observed in whole tissue and that realized within individual oligodendrocytes was expected, and a similar relationship likely exists in the PNS for Schwann cells (51). Samples were obtained at P7, a stage of maturation when significant myelin elaboration in cervical spinal cord has occurred (1); at P14, when myelin acquisition nears peak levels; at P21, when the levels of myelin protein mRNAs have declined from peak levels; at P30, when mature myelin maintaining cells predominate; and at P90, when mice are fully mature.

#### Wild type (WT)

*Mbp* mRNA was readily detectable at P7 and rapidly increased over the next week. Relative to P14, levels decreased to 85% by P21, 74% by P30 and 33% by P90; a developmental program consistent with prior investigations of multiple myelin gene expression programs and *Mbp* regulated reporters (1, 2, 41–43, 46, 52) (Fig. 2, Supplementary Table S1).

**Figure 2.**
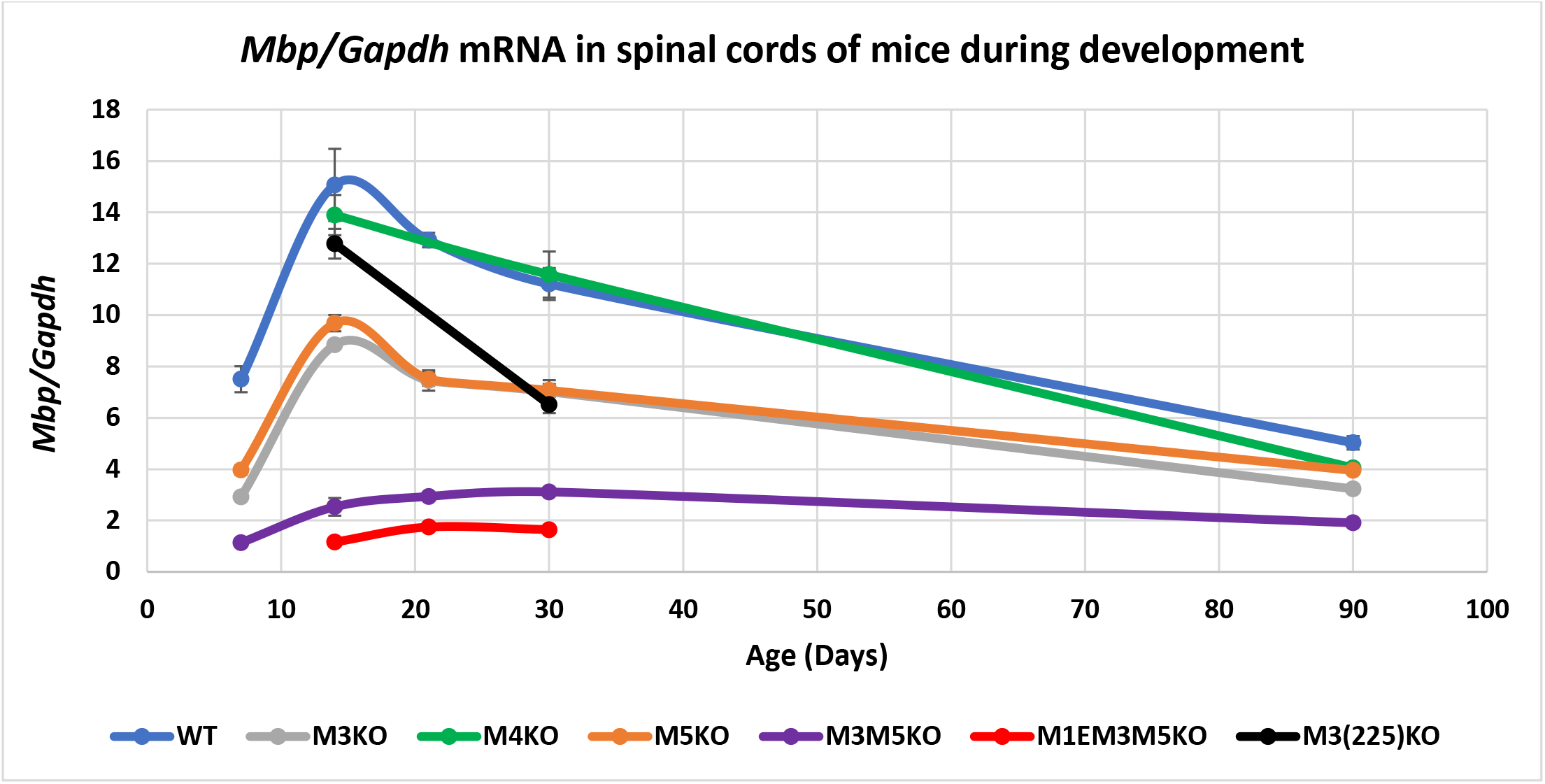
Relative *Mbp* mRNA accumulation in spinal cord. M1E, M3 and M5 are major enhancers of *Mbp* in oligodendrocytes. X-axis = post-natal age in days and Y-axis = *Mbp/Gapdh* mRNA ratio. Dots represent values at P7, 14, 21, 30 and 90 and connecting lines are the predicted developmental programs calculated as trendlines (Excel). Error bars = SEM

#### M5 KO lines

M5 function has not been investigated in transgenic preparations but its capacity to drive transcription in the oligodendrocyte CG-4 cell line was detected using transfected reporter constructs. The M5 enhancer displays limited interspecies conservation and contains a Myrf Chip-Seq binding profile that extends for 2466 bp encompassing a repeat domain (44). To establish if all M5 function is contained within the conserved non-repeat sequence, three M5 deletions were generated; the 829 bp deletion in the M3M5KO and M1EM3M5KO alleles removed only the conserved non-repeat sequence; the 1014 bp deletion (M5KOF6) extends 185 bp further 3’ to include a short repeat region; and the 3647 bp deletion (M5KO) encompasses the conserved 829 bp as well as 4 flanking repeat domains. Similar reductions in *Mbp* mRNA accumulation were observed with the 1014 and 3647 bp deletions thus demonstrating that the multiple repeat sequences do not influence M5 function (Supplementary Table S1).

#### M3KO, M3(225)KO, M5KO and M3M5 double KO

Oligodendrocytes in mice bearing either M3KO or M5KO alleles accumulated *Mbp* mRNA in similar developmental programs that differed quantitatively only at P7 and P90 when the M5KO allele supported relatively higher accumulation (Fig. 2, Supplementary Table S1). Notably, M3 and M5 differ in TF binding profiles including a binding peak for a master regulator, Myrf (44). The combined M3M5KO allele resulted in an additive reduction in *Mbp* mRNA accumulation at all ages with only a modest difference observed at P7. In addition, while M3M5KO mice demonstrated upregulation between P7 and P14 (typical of WT, M3KO and M5KO mice), the subsequent rapid downregulation by P21 was not observed, although some drop off occurred by P90. Consistent with the MBP being a limiting factor for myelin production, its reduced mRNA in M3M5KO mice resulted in a pronounced hypomyelination (Supplementary Figure S1). Unexpectedly, the partially truncated M3(225)KO allele that supported a strong and maintained up-regulation of reporter activity (43) led to equivalent or even modestly reduced *Mbp* mRNA accumulation at relative to M3 at ages examined.

#### M1EM3M5 triple KO

The *Mbp* proximal promoter sequence extending to −300 bp 5’ of the transcription start site failed to drive reporter expression in oligodendrocytes (46). However, a sequence extending 5’ to −377 bp (M1) supported transient expression during primary myelination thus demonstrating that the 5’ 77 bp (referred to as the M1 enhancer, M1E) is required for oligodendrocyte-specific activity. The M1EM3M5 allele investigated here is deleted of M3 and M5 as well as this enhancer-bearing 77 bp and the functional significance of the deleted 77 bp sequence is inferred from comparison of the M1EM3M5KO and M3M5KO alleles. M1EM3M5KO mice demonstrated a developmental expression program that paralleled that of M3M5KO mice but at approximately half the absolute level (Fig. 2, Supplementary Table S1). Post-weaning M1EM3M5KO mice accumulate Mbp mRNA to approximately 15% of WT and have a shivering phenotype similar to that of transgenic mice accumulating *Mbp* mRNA to 13.5% of WT (14).

#### M4KO

M4 drives reporter expression only in Schwann cells and consistent with that, the allele deleted of the M4 had no effect on *Mbp* mRNA accumulation in oligodendrocytes at P14 or P30. Surprisingly, moderately reduced accumulation was observed at P90 (Fig. 2, Supplementary Table S1).

### Mbp mRNA accumulation in sciatic nerve Schwann cells

#### WT and M4KO

In sciatic nerve samples from WT mice, *Mbp* mRNA was readily detectable at P4, reached a peak level at P14 and declined to 42% of the peak level by P90. Previously in reporter-based investigations, *Mbp* 5’ sequence extending to −8.9 kb (thus encompassing M1-M3) drove expression in oligodendrocytes but remained silent in Schwann cells. In contrast, the 5’ sequence extending to −9.4 kb, and thus incorporating the full M4 conserved sequence, expressed robustly in both oligodendrocytes and Schwann cells (42, 46). Consistent with the strong Schwann cell specific programming observed with all M4 bearing reporter constructs, M4KO mice accumulated *Mbp* mRNA at much reduced levels approximating 20% of WT at all ages (Fig. 3, Supplementary Table S2).

**Figure 3.**
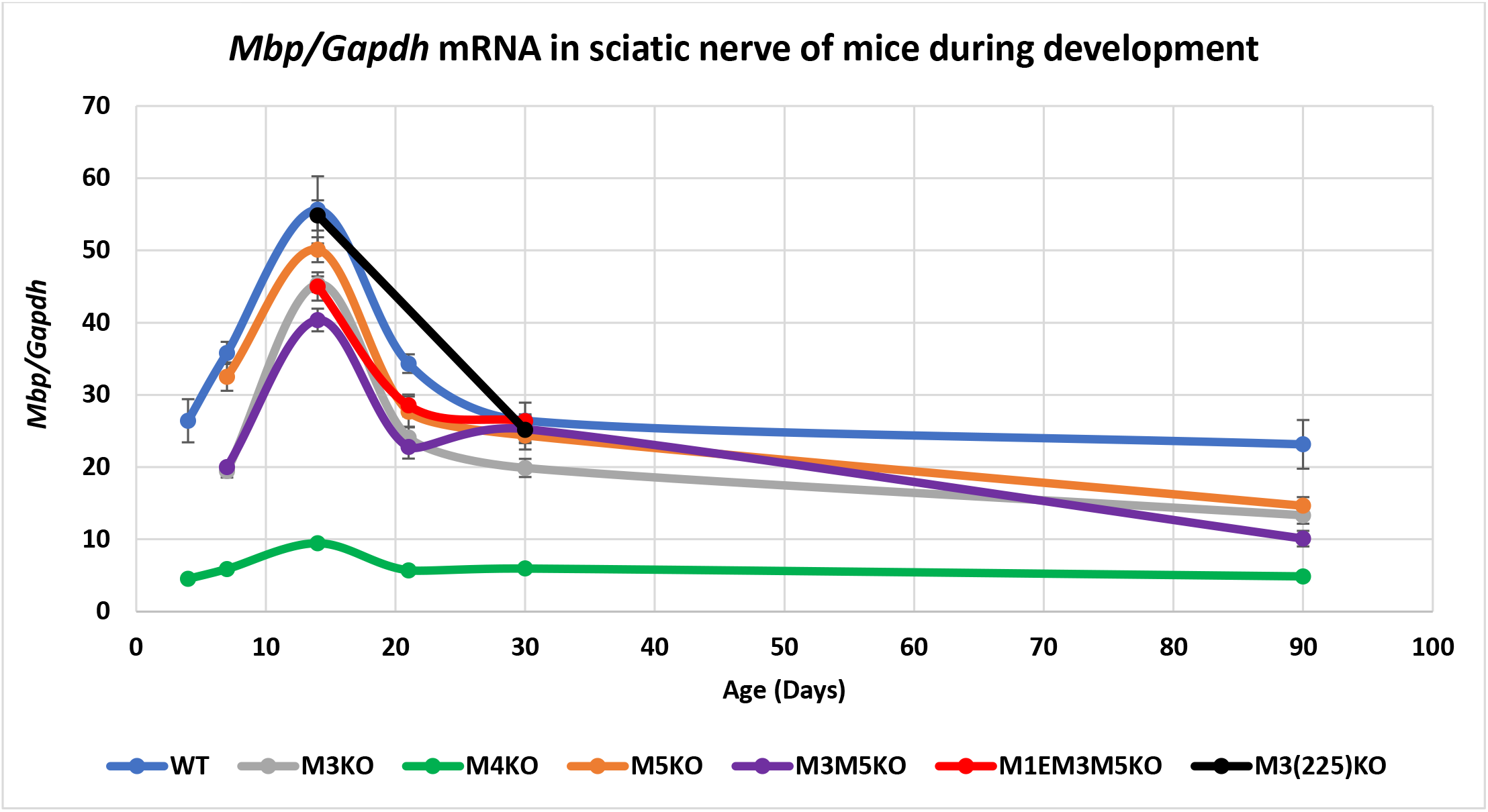
Relative *Mbp* mRNA accumulation in sciatic nerves. M4 is the major *Mbp* enhancer in Schwann cells. M3 and M5 also contribute while loss of M1E has no additional affect. X-axis = age in days and Y-axis = *Mbp/Gapdh* mRNA ratio. Dots represent post-natal days P4, 7, 14, 21, 30 and 90 and connecting lines are the predicted developmental programs calculated as trendlines (Excel). Error bars = SEM.

#### M3KO, M3(225)KO, M5KO, M3M5KO and M1EM3M5KO

In contrast to all M4 bearing reporter constructs, no M3 constructs driven through the *Mbp* proximal promoter (M1) expressed in Schwann cells. However, when M3 was ligated to a heterologous and minimal *hsp* promoter, Schwann cell expression was observed, albeit transiently during the preweaning period of active myelination (46). Thus, while M3 is not capable of productively engaging the *Mbp* proximal promoter in Schwann cells, it nonetheless binds relevant TFs during the period of active myelin synthesis in the PNS. All lines bearing the M3 deletion demonstrated reduced *Mbp* mRNA accumulation in Schwann cells not only preweaning but throughout development. In addition, while M5 has not been investigated for autonomous Schwann cell activity, the M5KO allele also led to reduced *Mbp* mRNA accumulation. Further, the reduction seen in mice bearing the combined M3M5KO allele trended lower than either the M3KO or M5KO alleles at most ages and was strictly additive at P14. Schwann cells in M1EM3M5KO mice also revealed an accumulation program comparable to that observed in M3M5KO mice while the partially deleted M3(225)KO allele had no effect on *Mbp* mRNA accumulation in Schwann cells (Fig. 3, Supplementary Table S2).

### Golli mRNA accumulation

In the mouse, *Golli* transcription initiates approximately 80 kb upstream of *Mbp*, and *Golli* isoforms variously incorporate *Mbp* exons as well as 264 bp of the *Mbp* proximal promoter (16, 53). The *Mbp* regulatory modules so far examined are located in the 20 kb 5’ of the *Mbp* start site and therefore within a *Golli* intron. In the oligodendrocyte lineage *Golli* accumulates predominantly in progenitors and is observed only rarely in mature myelinating oligodendrocytes (54). However, *Golli* is expressed by numerous cell types both within and beyond the nervous system including neurons and T-cells where distinct lineage specific developmental programs are observed (16, 54–58). Therefore, the *Golli* mRNA values obtained from spinal cord could arise from any combination of cell types. To determine levels of expression in nervous tissue devoid of oligodendrocytes we examined retina and for T-cells we examined thymus. Expression in Schwann cells was not determined as *Golli* accumulation was at the limit of detectability in sciatic nerve samples. Consistent with previous findings with optic nerve samples (42), *Golli* mRNA accumulation was reduced to 10% of WT in all tissues from lines in which M3 was deleted. In contrast, M5KO and M4KO lines demonstrated normal *Golli* mRNA accumulation in all tissues examined. Also, absence of the M1E sequence in the M1EM3M5KO allele had no further effect. Notably, in samples from M3(225)KO mice, *Golli* accumulation was normal at P14 indicating that the M3 subsequence deleted in this allele plays no role in *Golli* regulation at this age. However, at P30 *Golli* accumulation was 141% and 130% of WT in spinal cord and thymus respectively indicating an age-specific putative repressor role for the deleted sequence (Fig. 4 and 5, Supplementary Table S3).

**Figure 4.**
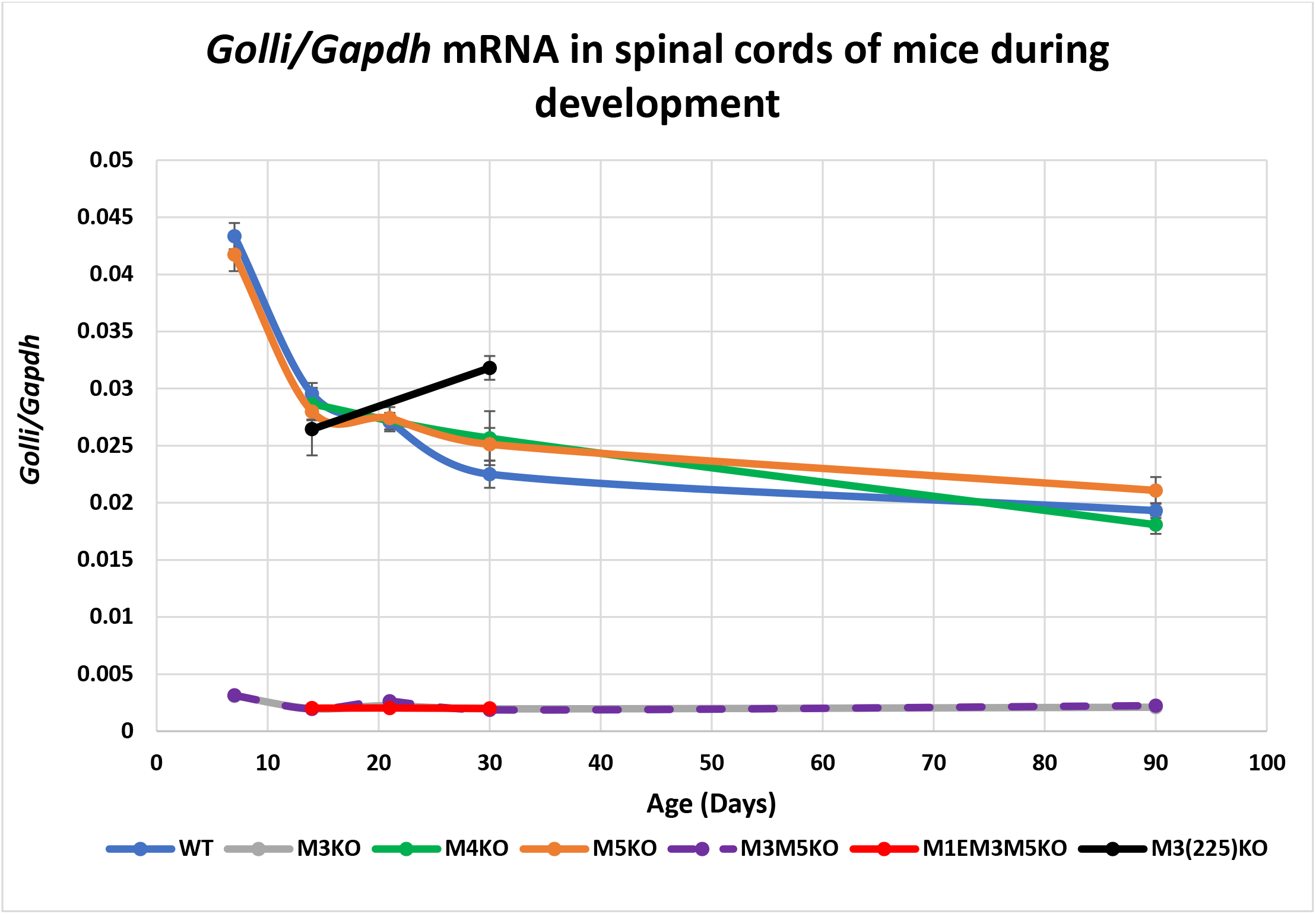
M3 is the major *Golli* enhancer in spinal cord. All alleles deleted of M3 demonstrate reduced *Golli* accumulation. In contrast, the partially deleted M3(225)KO allele did not reduce expression and at older ages even increased *Golli* accumulation. X-axis = age in days and Y-axis = *Golli/Gapdh* mRNA ratio. Dots represent post-natal days P7, 14, 21, 30 and 90 and connecting lines (both solid and dashed) are the predicted developmental programs calculated as trendlines (Excel). Error bars = SEM.

**Figure 5.**
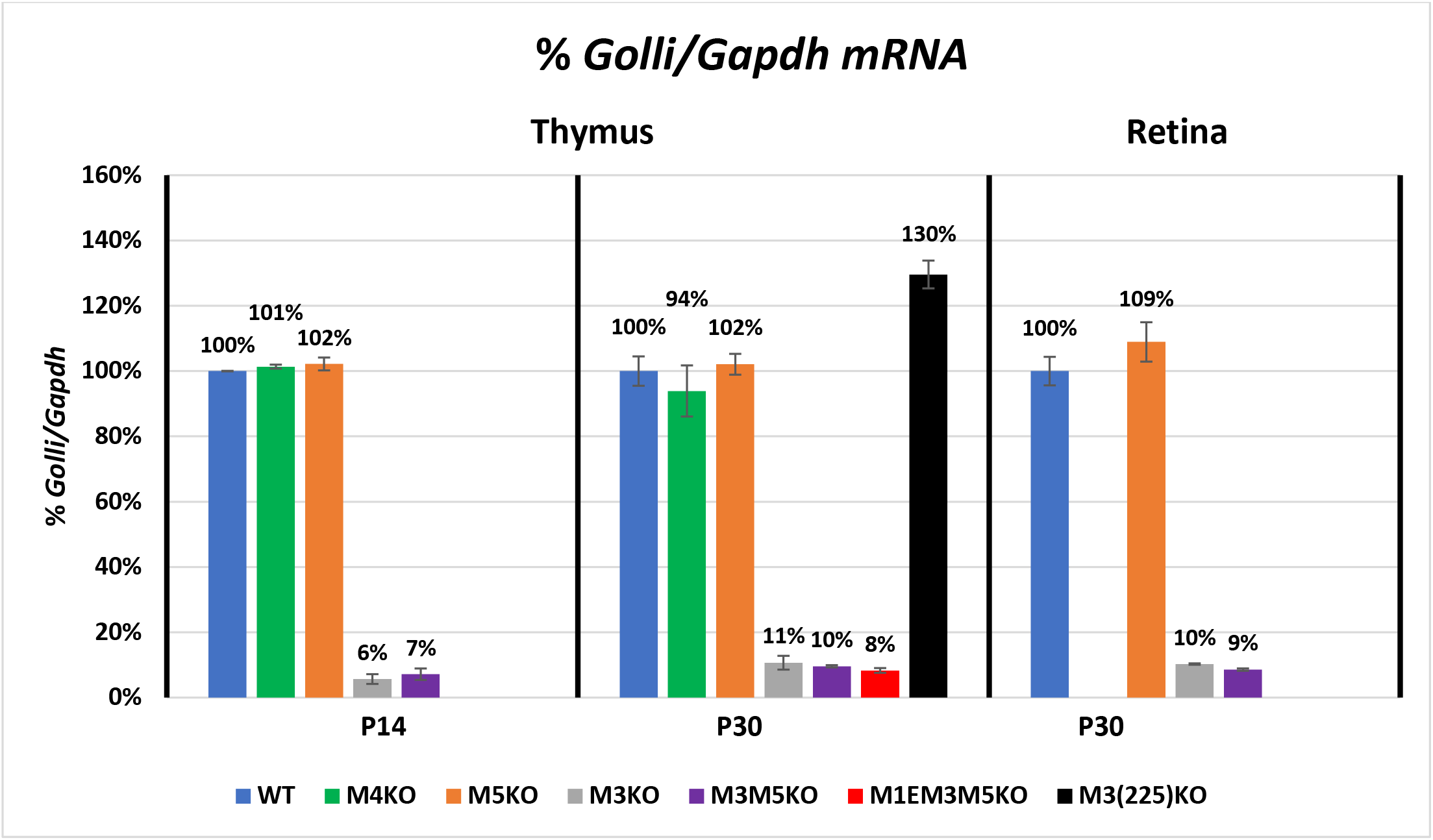
M3 is the major *Golli* enhancer in thymus and retina. All alleles deleted of M3, but not the partially deleted M3(225)KO allele, reduced *Golli* accumulation to a similar extent. X-axis = age in days and Y-axis = %*Golli/Gapdh* mRNA. Error bars = SEM.

## DISCUSSION

### Reporter genes vs enhancer KOs

Levels of chromatin organization beyond that associated with individual enhancers are thought to contribute to transcriptional regulation. Consequently, reporter constructs bearing isolated enhancers may not reveal the full extent of their regulatory activity. One such higher order level of organization is observed in super-enhancer domains that are enriched in active chromatin modifications and contain one or more transcriptionally active enhancers. They bind master TFs, mediator and co-activators and their extent is often demarcated by duplicated CTCF sites. Association between CTCF binding factor and cohesion supports the formation of chromatin loops creating insulated enhancer bearing neighborhoods (59). Additionally, such super-enhancers are frequently associated with key lineage-specific or lineage specifying genes (47, 59–61).

Multiple data sets from human brain revealed a super-enhancer domain extending through and 35 kb upstream of the *Mbp* gene (45, 60). Mouse cortical samples revealed a similar domain of 24 kb enriched in active transcription marks such as H3K27Ac and H3K4Me1 and demarcated by paired CTCF binding sites within which the M1-M5 enhancers are located (48, 62, 63). Consistently, in the CNS, M1, M3 and M5 (but not M4) demonstrate H3K27Ac enrichment and are bound by SOX10, a glial specifying transcriptional activator with a potential role in regulating the formation of super-enhancer domains (45). Further, SOX10 has been shown to interact with mediator in both oligodendrocytes and Schwann cells (29). Whether this super-enhancer domain forms exclusively in *Mbp*-expressing oligodendrocytes or also in Schwann cells and in the diverse *Golli*-expressing lineages remains to be determined. Nonetheless, in Schwann cells the M1, M3, M4 and M5 enhancers are all enriched in H3K27Ac and bind SOX10 (34, 45).

The extent to which super-enhancers confer synergy on their constituent enhancers remains controversial (60, 63–69). If inter-enhancer interactions occur within the *Golli/Mbp* super-enhancer domain, we expected them to be revealed through comparisons between the outputs of alleles bearing single and combined enhancer deletions. In oligodendrocytes, the double M3M5KO allele gave rise to the precise reduction predicted by the individual consequences of M3 and M5 deletions. Thus, no synergism between these 2 enhancers was detected in this lineage; a result consistent with the additive model in which super-enhancer activity equals the sum of its constituent enhancer activities.

The M4 enhancer revealed strong and autonomous targeting activity only Schwann cell in multiple reporter configurations (2, 41, 42, 46) and, as predicted, the M4KO allele resulted in a major (>80%) reduction in *Mbp* mRNA accumulation in Schwann cells. In contrast, all contiguous 5’ sequences terminating before M4 and encompassing M3 failed to drive reporter expression in Schwann cells, yet absence of M3 in KO lines caused a significant reduction of *Mbp* mRNA in Schwann cells. We identify this additional cryptic activity arising from the otherwise autonomous oligodendrocyte specific M3 enhancer as “stealth” activity. Defined in this manner, stealth activity in oligodendrocytes also originates from the autonomous lineage specific M4 Schwann cell enhancer as mRNA accumulation is reduced in the oligodendrocytes of M4KO mice, albeit only at P90. M5, shown to drive reporter expression in an oligodendrocyte cell line, may demonstrate the same capacity but its full autonomous targeting capacity in vivo is not yet known.

Few previous experiments have been designed to reveal such stealth enhancer activities assigned here to M3, M4 and potentially M5. It remains to be determined if this phenomenon is widespread or unique to *Golli/Mbp*, limited to loci expressed in multiple lineages or to enhancers within a common super-enhancer domain. However, that certain enhancers affect activity only through association with special “hub” enhancers has experimental support (65, 70) and the extent to which the stealth activity reported here is accommodated in that model remains to be determined. Specifically, M3 and M5 might participate in Schwann cells as non-hub enhancers engaging with M4 while in oligodendrocytes, M4 might act as a non-hub enhancer engaging with any of the oligodendrocyte enhancers. Unique to the previously hidden non-autonomous stealth activity observed here is its origin from otherwise fully autonomous enhancers that drive expression in an entirely different lineage.

### Reporter genes vs partial enhancer module KOs

Insight into the location of sequences engaged in proposed promoter-enhancer interactions was obtained from alleles bearing partial enhancer deletions. Absence of the 77 bp M1E sequence in the M1EM3M5KO allele reduced *Mbp* mRNA accumulation in oligodendrocytes beyond that imposed by the M3M5KO allele alone, consistent with an enhancing role for the deleted M1E domain in oligodendrocytes (Fig. 2). However, loss of M1E had no effect on *Mbp* mRNA accumulation in Schwann cells demonstrating that it has no Schwann cell enhancing activity nor a role in the productive engagement of M4 with the promoter components that remain in the M1EM3M5KO allele (Fig. 3).

A striking difference between the output of an enhancer KO allele and that predicted by a reporter gene was encountered with the M3(225) sub-sequence. This sequence drove reporter expression in oligodendrocytes approximately 5-fold higher than the intact M3 and when ligated directly to a minimal *hsp* promoter, it converted the transient pre-weaning activity of intact M3 in Schwann cells to constitutive expression at levels up to 50 fold higher (43). However, in the context of the endogenous locus, the same M3(225) truncation paradoxically had no effect in Schwann cells and reduced, rather than enhanced, *Mbp* mRNA accumulation in oligodendrocytes (Fig. 2 and 3). The basis for this striking difference between reporter construct activity compared to the same deletion in the endogenous locus remains unknown but the experimentally imposed close proximity of M3(225) and the *hsp* promoter in reporter constructs, may underlie the observed upregulation.

Further demonstration that distinct sequence sub-domains affect promoter-enhancer relationships was revealed by *Golli* expression. All alleles deleted of the intact M3 reduced *Golli* mRNA accumulation to approximately 10% of WT in all tissues at all ages examined (Fig. 4 and 5). In marked contrast, the partially truncated M3(225)KO allele supported near normal *Golli* accumulation at P14 with accumulation even modestly upregulated at P30 in both spinal cord and thymus. Thus, M3 sub-sequences differ in their capacity to engage the *Mbp* and *Golli* promoters, consistent with a model in which M3 functions through different TFs as a general “house-keeping” enhancer for *Golli* and a strong lineage-specific enhancer for *Mb*p.

### A model accommodating enhancer targeting activities

Here, the documented *Golli/Mbp* enhancer activities and their functional interactions lead to a DNA-looping model compatible with much of the integrated output of the locus (Fig. 6A and B). This model accommodates previously described differential TF binding (32, 59, 64, 71–75) and suggests that lineage-specific partnerships occur within sub-domains created by chromatin loops. Specifically, self-associating TF dimers, such as those formed by YY1 and SP1, are implicated in chromatin looping to support long-distance enhancer-promoter interaction (49, 76–79). This model can also accommodate the emergence of lineage specific regulatory programs such that the output of a single locus can evolve to match the unique requirements of the different myelinating cells in the CNS and PNS. However, this model does not take into account the mediators of stealth activity.

**Figure 6.**
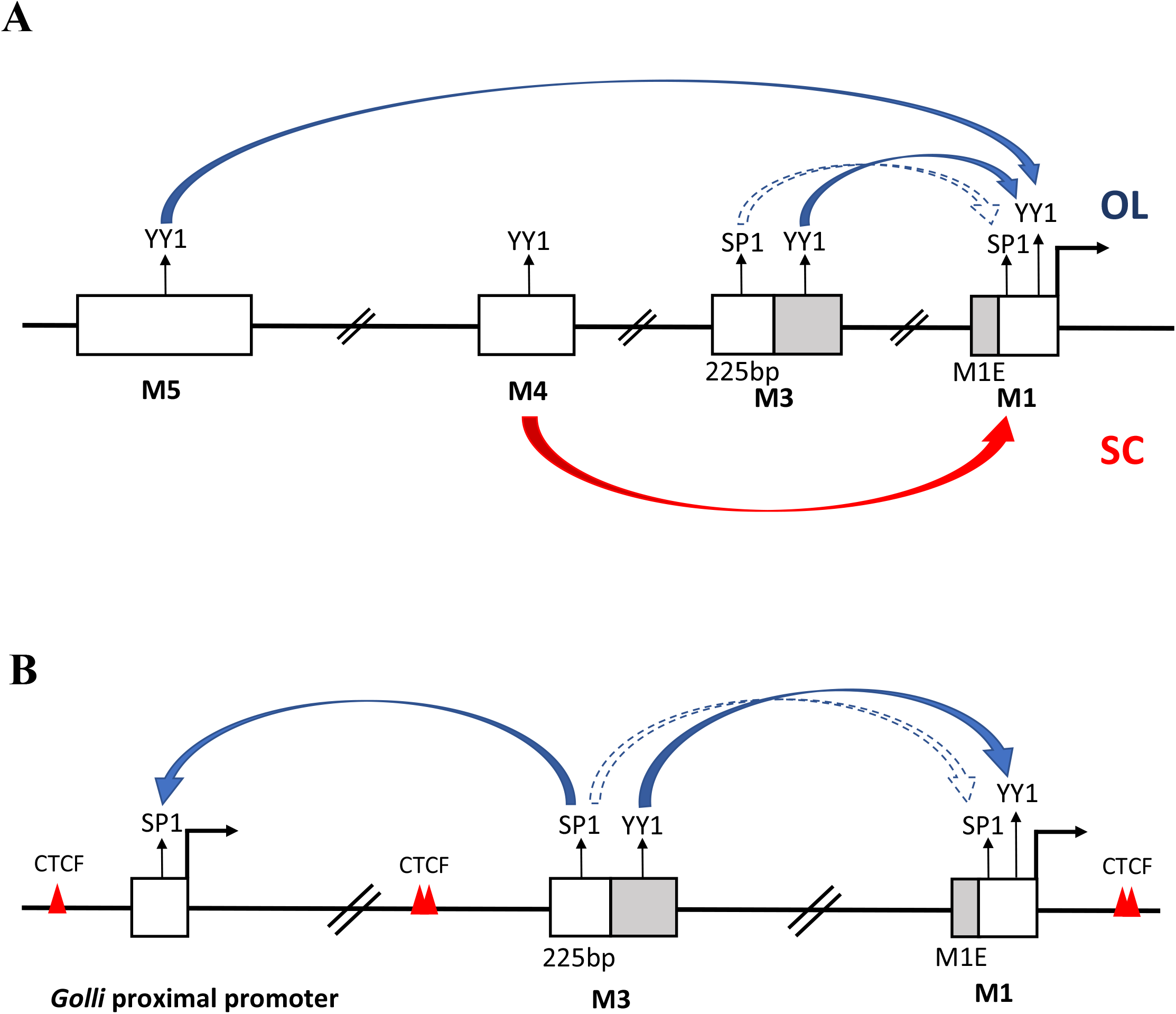
A tentative DNA looping model for enhancer engagement. (**A**) The predicted direct enhancer-promoter interaction for *Mbp* expression in oligodendrocytes (top with blue arrows) and Schwann cells (bellow with the red arrow). (**B**) The predicted relationship between M3 and the *Mbp* and *Golli* promoters. White boxes = enhancers and proximal promoters. Deleted sequences are highlighted in grey. Filled arrows = continuous potential interactions. Dotted arrows = potential interactions occurring at young age. Red triangles = CTCF binding sites.

M1, M3, M4, M5 and the *Golli* proximal promoter all have predicted YY1 binding sites while only M3, M3(225) and the *Golli* proximal promoter have SP1 binding sites (Jaspar; relative score (rs) > 90%) (80), however lower affinity SP1 and YY1 motifs (80% < rs < 90%) exist in all. Accordingly, in oligodendrocytes, M3 and M5 interaction with the promoter could involve YY1 dimerization (Fig. 6A) and consistent with this, the M3(225)KO allele, lacking the sequence containing YY1 binding site, resulted in the same *Mbp* mRNA reduction as observed in M3KO mice (P30). Additionally, conditional KO of YY1 led to amyelination and behavioral phenotypes characteristic of MBP null shiverer mice (81, 82). However, during active myelination at P14, the M3(225)KO allele conferred a more modest reduction than the M3KO allele. During with this period, SP1 accumulation transiently increases and becomes phosphorylated via the PKC/Erk pathway (83, 84). As the SP1 binding site remains in the M3(225)KO allele and the SP1 phosphorylated through that pathways has been shown to increase its DNA binding capacity (85), the partial restoration of enhancer-promoter interaction in the absence of the YY1 binding site could be conferred through SP1 (Fig. 6A).

Although enhancer activity mediated largely by YY1 dimerization conveniently accommodates our observations on *Mbp* expression, it fails to account for *Golli* programming (Fig. 6B). While the *Golli* proximal promoter contains both YY1 and SP1 binding motifs, among the enhancers investigated here, only the oligodendrocyte M3 enhancer that contains an SP1 binding site modulates *Golli* output (80). Consistent with this model, *Golli* expression driven by the truncated enhancer in M3(225)KO mice was anticipated by its retained SP1 binding site and hence its predicted capacity to associate with the *Golli* promoter (Fig. 6B).

### Enhancer activity during development

Beyond insight into the functional organization of *Golli/Mbp* regulatory sequences, the developmental programs conferred by *Mbp* enhancer-regulated reporter genes and KO alleles illuminate further aspects of oligodendrocyte biology. As demonstrated by the capacity of oligodendrocytes to myelinate inanimate fibers *in vitro*, initial myelin elaboration can be supported entirely by oligodendrocyte intrinsic programming (86). However, it is now widely recognized that myelin in the mature CNS can demonstrate plastic changes potentially in response to neuronal activity (3, 4, 87, 88). Here, the M3(225)KO allele showed differential age-related regulatory capacity for both *Golli* and *Mbp* in oligodendrocytes (P14 vs P30). Similarly, the M4KO allele differentially affected *Mbp* mRNA accumulation in oligodendrocytes at P14 and P30 vs P90. These observations are consistent with a model in which an evolving transcription factor repertoire engages these enhancers at different stages of maturation. Conversely, in the absence of M3 and M5 (M3M5KO and M1EM3M5KO) the enhancer activity remaining is not sufficient to elicit the normal preweaning peak of *Mbp* mRNA accumulation in oligodendrocytes thus demonstrating that developmentally contextual activity is not distributed equivalently among all regulatory components.

Our observations provide a functional framework from which the role of chromatin configuration, modification and specific TF binding across the *Golli/Mbp* locus can be approached (65). The mouse models described here contain unique configurations of *Golli/Mbp* regulatory sequence and these differences may facilitate definition of the conditions that foster the formation of super-enhancers and help expose their functional significance. Finally, these enhancer-deleted mouse models elaborate myelin sheathes with variably reduced thickness providing novel opportunities to illuminate basic features of the axon-myelin relationship.

## MATERIAL AND METHODS

### Animals

All experiments were carried out in accordance with the guidelines of the Canadian Council on Animal Care and the McGill University Animal Care Committee.

### CRISPR design and gene editing

#### M4 and M5 sequences

The 422bp M4 enhancer targeted here was described previously (41). M5 refers to the target of the MYRF ChIP carried out in rat (44). Using the UCSC browser, this sequence was aligned with the mouse genome (chr18:82,536,728-82,539,249 GRCm38/mm10). For the purpose of generating M4 and M5 enhancer knock-outs (KOs), single guide RNAs (sgRNAs) were designed to target sequences flanking the conserved enhancer domain such that double strand breaks would be simultaneously introduced at both sides of the enhancer resulting in deletion of the intervening enhancer sequence. sgRNAs were designed (using the CRISPR Design http://crispr.mit.edu/) (89) to identify locus specific targets. To minimize the potential impact of inefficient sgRNAs, we designed 2 that bind in close proximity (for a total of 4 sgRNAs per enhancer deletion). The sgRNA target sequences used to generate the KO mice are indicated in Supplementary Table S4. For the M5 and M4 deletions, all sgRNAs were apparently effective.

#### sgRNA design

The plasmid DR274 was a gift from Keith Joung (Addgene plasmid # 42250) (90). DR274 was digested with BsaI which cuts twice between the T7 promoter and the gRNA scaffold, leaving sticky ends. For each of the targets listed above, two oligos, one for each strand, were ordered from IDT (Supplementary Table S5). They were annealed at 40uM each in NEB3 buffer. Each has one of the DR274 sticky ends so that they could be ligated into the plasmid using the NEB Quick Ligation Kit (M2200S). Each ligation mix was transformed into competent bacteria and kanamycin resistant clones obtained. 3 clones of each were sequenced in the relevant region and used to generate the sgRNA. To generate the sgRNA template for the M4 deletion, the PCR method using two long overlapping oligos was used (Supplementary Table S5) (91). The MEGA shortscript T7 kit from Life Technologies was used to synthesize the sgRNA from the T7 promoter. The resulting sgRNA was tested for integrity on a Bioanlyzer at the McGill Genome Center.

The target-specific crRNAs (Supplementary Table S5) were hybridized to Alt-R^®^ CRISPR-Cas9 tracrRNA, the Universal 67mer tracrRNA from IDT to generate the functional sgRNAs according to the manufacturer’s instructions. AltR1 and AltR2 are proprietary (IDT) modifications to increase the stability of these short RNAs.

#### Zygote manipulation, delivery of CRISPR components and transplantation into pseudopregnant mice

Zygotes were recovered mid-day from the oviduct of WT or M3KO C57Bl/6 mice (42) naturally mated to wmN2 transgenic mice (92). The cumulus cells were removed by a short incubation in 1% hyaluronidase/M2 medium (Millipore) and moved into Advanced KSOM media (Millipore) at 37°C with 5% CO2. All zygote manipulation was done at room temperature and the media was kept under mineral oil. M4KO mice were generated by cytoplasmic microinjection of 25ng/ul Cas9 mRNA (PNA Bio) and 12.5ng/ul of each of four sgRNAs. M5KO mice were generated by electroporation. Prior to electroporation the zygotes were moved to Opti-MEM (Life Technologies) and thinning of the zona was achieved by treating the zygotes with Acid Tyrode’s solution (Millipore) for 10 seconds and transferring them back into fresh Opti-MEM. Zygotes were electroporated according to the ZEN2 protocol (93, 94) with a final concentration of 250ng/ul Cas9 mRNA (PNA Bio) and 300ng/ul sgRNA dissolved in TE pH7.5/Opti-MEM at a 1:1 ratio (93). A 20ul drop of this mix containing the CRISPR reagents was prepared and the batch of 30-50 zygotes carried in less than 1ul of Opti-MEM were moved into this drop. The mix was transferred to a 1 mm electroporation cuvette purchased from BioRad and electroporation was carried out using a Bio-Rad Gene Pulser Xcell electroporator. Embryos were subjected to 1-2 pulses of 25-30 V according to the ZEN protocol (93). After microinjection or electroporation embryos were cultured overnight in advanced KSOM media at 37°C with 5% CO2. After overnight incubation, embryos at the 2-cell stage were transplanted (bilaterally, approximately 15/mouse) into the fallopian tubes of CD1 female recipients rendered pseudopregnant by mating with B6C3F1 vasectomized males (purchased from Charles River).

#### Genotyping and breeding scheme

Pups were tail-biopsied at weaning for genotyping. Tail samples were digested at 55C overnight in lysis buffer (containing 100 mM Tris, pH 8.0, 5 mM EDTA, pH8.0, 200 mM NaCl, 0.2% sodium dodecyl sulfate (SDS) and 100ug/ul proteinase K) and genomic DNA was extracted. Genotyping initially was done using PCR with primers surrounding the sequence to be deleted. Upon detection of a desired, shorter-than-WT, band, the PCR product was sequenced at the McGill University and Génome Québec Innovation Centre and the existence of M4 and M5 deletions confirmed. Founder mice were mated to WT C57Bl/6 and the consequent progeny were genotyped by PCR for the deletion and LacZ (to detect the presence of a transgene at the HPRT locus, that exists within our donor colony and select against it). Mice carrying the enhancer deletion were mated to homozygosity while breeding out the transgene located on the X chromosome. In total, 2 lines of M4KO mice (identical sequencing results), 3 lines of M5KO (bearing different deletion lengths) 1 line of M3M5KO double KO and 1 line of M1EM3M5KO triple KO, were established.

#### Tissue samples

After the homozygous lines of mice were established, samples from males and females of WT, M4KO, M3KO, M5KO and M3M5KO lines were taken at the ages indicated. The mice were anesthetized with a lethal dosage of Avertin and sciatic nerve, cervical spinal cord, retina and thymus samples were collected into RNAlater solution (Ambion) according to the manufacturer’s instructions and stored at −20°C.

#### RNA extraction and qRT-PCR

Total RNA extraction was done using Trizol (Life Technologies) and a Qiagen RNeasy MinElute Cleanup kit. RNA was eluted in nuclease free water and its concentration was measured using a spectrophotometer. The RT reaction was carried out using Superscript IV VILO Mastermix (Life Technologies) using 1ug of total RNA according to the manufacturer’s instructions and the resulting cDNA was stored at −80C. A QuantStudio™ 7 Flex Real-Time PCR System (Life technologies) was used for qPCR in a 96-well plate. On the day of qPCR, the cDNA was diluted 20x and 40x for measuring *Golli* and 100x and 400x for measuring *Mbp* and *Gapdh*. Each sample was measured twice at the low dilution and once at the high dilution. Samples from WT, M4KO, M5KO, M3KO, M3M5KO mice were measured on a single 96-well plate to avoid inter-plate variability. To measure *Mbp* and *Gapdh* in SN and cervical spinal cord, Taqman probes (*Mbp*: Mm01266402_m1, *Gapdh:* Mm99999915_g1, Life technologies) were used. For *Golli* measurements in cervical spinal cord and thymus however, the SYBR green method was used (PowerUp SYBR green master mix, Life Technologies). Multiple primer sets were designed, tested and the optimal pair (2F: 5’ATTGGGTCGCCATGGGAAAC, 2B: 5’CCAGCCTCTCCTCGGTGAAT) was chosen. On each plate, 5 10-fold serial dilutions of a DNA standard were run in triplicate to generate a standard curve. Standards were prepared by amplifying a sequence larger than the measured amplicon. After standard PCR, the single band was purified from a gel with a NucleoSpin Gel and PCR cleanup kit (Macherey-Nagel) and its concentration determined. The efficiencies of reactions for both Taqman and SYBR green methods inferred from standard curves were 95-105%.

#### Data analysis

After the generation of data, the measurements of each sample from both dilutions were averaged. Relative amounts of *Mbp* and *Golli* were calculated by dividing the average of each by the average of *Gapdh* for the same sample. The relative *Mbp* and *Golli* measurements of all samples of each mouse line were averaged and the standard deviation was calculated and normalized to measured WT levels. As analysis of sub-samples revealed no gender differences, male and female samples were combined throughout this investigation.

#### Light and electron microscopy

Mice lethally anesthetized with Avertin, were transcardially perfused with 2.5% glutaraldehyde + 0.5% paraformaldehyde in 0.1M sodium cacodylate buffer and cervical spinal cord and ON samples were collected. Samples were postfixed overnight at 4°C followed by rinsing with 0.1M sodium cacodylate buffer. A second post fixation was done with 1% osmium tetroxide followed by rinsing with ddH2O. Samples were dehydrated by incubation in increasing concentrations of acetone: 30%, 50%, 70%, 80%. 90% and 3X100%. Infiltration was done with 1:1, 2:1, 3:1 (epon:acetone) followed by embedding in epon and overnight polymerization at 6Ü°C. 0.5 um sections were stained with Toluidine blue and cover slips were mounted with epon for imaging. Slides were imaged with 63X or 100X oil immersion objectives by light microscopy (Zeiss Axio Imager M1).

## ACKNOWLEDGMENT

McGill facilities: Facility for Electron Microscopy Research (FEMR). McGill University and Genome Quebec Innovation Centre.

## AUTHOR CONTRIBUTIONS

H.B., H.F., A.P.: Designed and conducted experiments, wrote manuscript. K.S.: Designed experiments and reviewed manuscript.

## SUPPORTING INFORMATION LEGENDS

**Supplementary Figure S1.** Mice bearing the M3M5KO allele demonstrate CNS hypomyelination. Electron micrographs were obtained from cross sections of the ventral medial cervical spinal cord from P90 WT and M3M5 mice (**A** and **B** at 640x) and (**C** and **D** at 3000x). The axon population in this domain ranges from small to large calibers. In the M3M5KO sample, axons of all calibers are typically ensheathed with compact myelin markedly thinner than normal although rare small calibre axons lacking compact myelin (*) were encountered.

**Supplementary Table S1.** Relative *Mbp* mRNA analysis in spinal cord of enhancer knock-out mice at P7, P14, P21, P30 and P90. The values are presented as % ± standard error of the mean. “*” and “**” represent p-values ≤ 0.05 and ≤ 0.01 respectively. n(F:M) represents the number of Female and Male mice from each genotype analyzed at each age.

**Supplementary Table S2:** Relative *Mbp* mRNA analysis in sciatic nerve of enhancer knock-out mice at P4, P7, P14, P21, P30 and P90. The values are presented as % ± standard error of the mean. “*” and “**” represent p-values ≤ 0.05 and ≤ 0.01 respectively. n(F:M) represents the number of Female and Male mice from each genotype analyzed at each age. “#” indicates that sciatic nerves from two mice were combined for each sample.

**Supplementary Table S3:** Relative *Golli* mRNA accumulation in spinal cord of enhancer knock-out mice at P7, P14, P21, P30 and P90. The values are presented as % ± standard error of the mean. “**” represent p-values ≤ 0.01. n(F:M) represents number of Female and Male mice analyzed.

**Supplementary Table S4.** sgRNA target sequences used to generate the KO mice.

**Supplementary Table S5.** Oligonucleotides used for CRISPR editing.

## References

1. Foran DR, Peterson A. Myelin Acquisition in the Central Nervous System of the Mouse Revealed by an MBP-Lac Z Transgene. The Journal of Neuroscience. 1992;12:4890–7.

2. Forghani R, Garofalo L, Foran DR, Farhadi HF, Lepage P, Hudson TJ, et al. A Distal Upstream Enhancer from the Myelin Basic Protein Gene Regulated Expression in Myelin-Forming Schwann Cells. The Journal of Neuroscience. 2001;21(11):3780–7.

3. Bercury KK, Macklin WB. Dynamics and mechanisms of CNS myelination. Dev Cell. 2015;32(4):447–58.

4. Bergles DE, Richardson WD. Oligodendrocyte Development and Plasticity. Cold Spring Harb Perspect Biol. 2015;8(2).

5. Sock E, Wegner M. Transcriptional control of myelination and remyelination. Glia. 2019;67(11):2153–65.

6. Vassall KA, Bamm VV, Harauz G. MyelStones: the executive roles of myelin basic protein in myelin assembly and destabilization in multiple sclerosis. Biochem J. 2015;472(1):17–32.

7. Peterson AC, Bray GM. Hypomyelination in the peripheral nervous system of shiverer mice and in shiverer in equilibrium normal chimaera. The Journal of Comparative Neurology. 1984;227:348–56.

8. Rosenbluth J. Peripheral myelin in the mouse mutant Shiverer. The Journal of Comparative Neurology. 1980;193:729–39.

9. Gould RM, Byrd AL, Barbarese E. The number of Schmidt-Lanterman incisures is more than doubled in shiverer PNS myelin sheaths. Journal of Neurocytology. 1995;24:85–98.

10. Smith-Slatas C, Barbarese E. Myelin basic protein gene dosage effects in the PNS. Molecular and cellular neurosciences. 2000;15(4):343–54.

11. Bird T, Farrel, D.F. and Sumi, S.H.,. Genetic development myelin defect in Shiverer mouse. Neurochemistry. 1977;8:153.

12. Rosenbluth J. Central myelin in the mouse mutant Shiverer. The Journal of Comparative Neurology. 1980;194:639–48.

13. Chernoff G. Shiverer: an autosomal recessive mutant mouse with myelin deficiency. The journal of Heredity. 1981;72:128.

14. Popko B, Puckett C, Lai E, Shine H, Readhead C, Takahashi N, et al. Myelin deficient mice: expression of myelin basic protein and generation of mice with varying levels of myelin. Cell. 1987;48:713–21.

15. Shine HD, Readhead C, Popko B, Hood L, Sidman RL. Morphometric Analysis of Normal, Mutant, and Transgenic CNS: Correlation of Myelin Basic Protein Expression to Myelinogenesis. Journal of Neurochemistry. 1992;58(1):342–9.

16. Campagnoni A, Pribyl T, Campagnoni C, Kampf K, Amur-Umarjee S, Landry C, et al. Structure and developmental regulation of Golli-mbp, a 105-kilobase gene that encompasses the myelin basic protein gene and is expressed in cells in the oligodendrocyte lineage in the brain. Biological chemistry. 1993;268:4930–8.

17. Feng JM, Fernandes AO, Campagnoni CW, Hu YH, Campagnoni AT. The golli-myelin basic protein negatively regulates signal transduction in T lymphocytes. J Neuroimmunol. 2004;152(1-2):57–66.

18. Feng JM, Givogri IM, Bongarzone ER, Campagnoni C, Jacobs E, Handley VW, et al. Thymocytes Express the golli Products of the Myelin Basic Protein Gene and Levels of Expression Are Stage Dependent. The Journal of Immunology. 2000;165(10):5443–50.

19. Feng JM, Hu YK, Xie LH, Colwell CS, Shao XM, Sun XP, et al. Golli protein negatively regulates store depletion-induced calcium influx in T cells. Immunity. 2006;24(6):717–27.

20. Paez PM, Cheli VT, Ghiani CA, Spreuer V, Handley VW, Campagnoni AT. Golli myelin basic proteins stimulate oligodendrocyte progenitor cell proliferation and differentiation in remyelinating adult mouse brain. Glia. 2012;60(7):1078–93.

21. Paez PM, Fulton DJ, Spreuer V, Handley V, Campagnoni CW, Campagnoni AT. Regulation of store-operated and voltage-operated Ca2+ channels in the proliferation and death of oligodendrocyte precursor cells by golli proteins. ASN Neuro. 2009;1(1).

22. Vt C, Da SG, V S, V H, At C, Pm P. Golli Myelin Basic Proteins Modulate Voltage-Operated Ca(++) Influx and Development in Cortical and Hippocampal Neurons. Mol Neurobiol. 2016;53(8):5749–71.

23. Berndt JA, Kim JG, Tosic M, Kim C, Hudson LD. The transcriptional regulator Yin Yang 1 activates the myelin PLP gene. Journal of Neurochemistry. 2001;77(3):935–42.

24. Wang SZ, Dulin J, Wu H, Hurlock E, Lee SE, Jansson K, et al. An oligodendrocyte-specific zinc-finger transcription regulator cooperates with Olig2 to promote oligodendrocyte differentiation. Development. 2006;133(17):3389–98.

25. Cahoy JD, Emery B, Kaushal A, Foo LC, Zamanian JL, Christopherson KS, et al. A transcriptome database for astrocytes, neurons, and oligodendrocytes: a new resource for understanding brain development and function. The Journal of neuroscience: the official journal of the Society for Neuroscience. 2008;28(1):264–78.

26. Svaren J, Meijer D. The molecular machinery of myelin gene transcription in Schwann cells. Glia. 2008;56(14): 1541–51.

27. Emery B, Agalliu D, Cahoy JD, Watkins TA, Dugas JC, Mulinyawe SB, et al. Myelin gene regulatory factor is a critical transcriptional regulator required for CNS myelination. Cell. 2009;138(1):172–85.

28. Ahrendsen JT, Macklin W. Signaling mechanisms regulating myelination in the central nervous system. Neuroscience Bulletin. 2013;29(2):199–215.

29. Vogl MR, Reiprich S, Kuspert M, Kosian T, Schrewe H, Nave KA, et al. Sox10 cooperates with the mediator subunit 12 during terminal differentiation of myelinating glia. The Journal of neuroscience: the official journal of the Society for Neuroscience. 2013;33(15):6679–90.

30. Hung HA, Sun G, Keles S, Svaren J. Dynamic regulation of Schwann cell enhancers after peripheral nerve injury. The Journal of biological chemistry. 2015;290(11):6937–50.

31. Stolt CC, Wegner M. Schwann cells and their transcriptional network: Evolution of key regulators of peripheral myelination. Brain Res. 2015.

32. Weider M, Wegner M. SoxE factors: Transcriptional regulators of neural differentiation and nervous system development. Seminars in cell & developmental biology. 2017;63:35–42.

33. Jang SW, Srinivasan R, Jones EA, Sun G, Keles S, Krueger C, et al. Locus-wide identification of Egr2/Krox20 regulatory targets in myelin genes. J Neurochem. 2010;115(6):1409–20.

34. Srinivasan R, Sun G, Keles S, Jones EA, Jang SW, Krueger C, et al. Genome-wide analysis of EGR2/SOX10 binding in myelinating peripheral nerve. Nucleic acids research. 2012;40(14):6449–60.

35. Chen Y, Wang H, Yoon SO, Xu X, Hottiger MO, Svaren J, et al. HDAC-mediated deacetylation of NF-kappaB is critical for Schwann cell myelination. Nature neuroscience. 2011;14(4):437–41.

36. Weider M, Reiprich S, Wegner M. Sox appeal - Sox10 attracts epigenetic and transcriptional regulators in myelinating glia. Biological chemistry. 2013;394(12):1583–93.

37. Emery B, Lu QR. Transcriptional and Epigenetic Regulation of Oligodendrocyte Development and Myelination in the Central Nervous System. Cold Spring Harb Perspect Biol. 2015;7(9):a020461.

38. Scaglione A, Patzig J, Liang J, Frawley R, Bok J, Mela A, et al. PRMT5-mediated regulation of developmental myelination. Nat Commun. 2018;9(1):2840.

39. Tiane A, Schepers M, Rombaut B, Hupperts R, Prickaerts J, Hellings N, et al. From OPC to Oligodendrocyte: An Epigenetic Journey. Cells. 2019;8(10).

40. The Encode Project Consortium. An integrated encyclopedia of DNA elements in the human genome. Nature. 2012;489(7414):57–74.

41. Denarier E, Forghani R, Farhadi HF, Dib S, Dionne N, Friedman HC, et al. Functional organization of a Schwann cell enhancer. The Journal of neuroscience: the official journal of the Society for Neuroscience. 2005;25(48): 11210–7.

42. Dib S, Denarier E, Dionne N, Beaudoin M, Friedman HH, Peterson AC. Regulatory modules function in a non-autonomous manner to control transcription of the mbp gene. Nucleic acids research. 2011;39(7):2548–58.

43. Dionne N, Dib S, Finsen B, Denarier E, Kuhlmann T, Drouin R, et al. Functional organization of an Mbp enhancer exposes striking transcriptional regulatory diversity within myelinating glia. Glia. 2016;64(1):175–94.

44. Bujalka H, Koenning M, Jackson S, Perreau VM, Pope B, Hay CM, et al. MYRF is a membrane-associated transcription factor that autoproteolytically cleaves to directly activate myelin genes. PLoS Biol. 2013;11(8):e1001625.

45. Lopez-Anido C, Sun G, Koenning M, Srinivasan R, Hung HA, Emery B, et al. Differential Sox10 genomic occupancy in myelinating glia. Glia. 2015.

46. Farhadi HF, Lepage P, Forghani R, Friedman HCH, Orfali W, Jasmin L, et al. A Combinatorial Network of Evolutionarily Conserved Myelin Basic Protein Regulatory Sequence Confers Distinct Gial-Specific Phenotypes. Neurosciene. 2003;23(32):10214–23.

47. Whyte WA, Orlando DA, Hnisz D, Abraham BJ, Lin CY, Kagey MH, et al. Master transcription factors and mediator establish super-enhancers at key cell identity genes. Cell. 2013;153(2):307–19.

48. Khan A, Zhang X. dbSUPER: a database of super-enhancers in mouse and human genome. Nucleic acids research. 2016;44(D1):D164–D71.

49. Nolis IK, McKay DJ, Mantouvalou E, Lomvardas S, Merika M, Thanos D. Transcription factors mediate long-range enhancer–promoter interactions. Proceedings of the National Academy of Sciences. 2009;106(48):20222–7.

50. Bu J, Banki A, Wu Q, Nishiyama A. Increased NG2(+) glial cell proliferation and oligodendrocyte generation in the hypomyelinating mutant shiverer. Glia. 2004;48(1):51–63.

51. Grove M, Kim H, Santerre M, Krupka AJ, Han SB, Zhai J, et al. YAP/TAZ initiate and maintain Schwann cell myelination. eLife. 2017;6.

52. Fulton DL, Denarier E, Friedman HC, Wasserman WW, Peterson AC. Towards resolving the transcription factor network controlling myelin gene expression. Nucleic acids research. 2011;39(18):7974–91.

53. Mathisen PM, Pease S, Garvey J, Hood L, Readhead C. Identification of an embryonic isoform of myelin basic protein that is expressed widely in the mouse embryo. Proceedings of the National Academy of Sciences. 1993;90(21):10125–9.

54. Filipovic R, Rakic S, Zecevic N. Expression of Golli proteins in adult human brain and multiple sclerosis lesions. Journal of Neuroimmunology. 2002;127(1):1–12.

55. Landry C, Ellison J, Pribyl T, Campagnoni C, Kampf K, Campagnoni A. Myelin basic protein gene expression in neurons: developmental and regional changes in protein targeting within neuronal nuclei, cell bodies, and processes. The Journal of Neuroscience. 1996;16(8):2452–62.

56. Landry CF, Ellison J, Skinner E, Campagnoni AT. Golli-MBP proteins mark the earliest stages of fiber extension and terminal arboration in the mouse peripheral nervous system. Journal of Neuroscience Research. 1997;50(2):265–71.

57. Landry CF, Pribyl TM, Ellison JA, Givogri MI, Kampf K, Campagnoni CW, et al. Embryonic Expression of the Myelin Basic Protein Gene: Identification of a Promoter Region That Targets Transgene Expression to Pioneer Neurons. The Journal of Neuroscience. 1998; 18(18):7315–27.

58. Feng JM. Minireview: expression and function of golli protein in immune system. Neurochem Res. 2007;32(2):273–8.

59. Dowen JM, Fan ZP, Hnisz D, Ren G, Abraham BJ, Zhang LN, et al. Control of cell identity genes occurs in insulated neighborhoods in mammalian chromosomes. Cell. 2014;159(2):374–87.

60. Hnisz D, Abraham BJ, Lee TI, Lau A, Saint-Andre V, Sigova AA, et al. Super-enhancers in the control of cell identity and disease. Cell. 2013;155(4):934–47.

61. Pott S, Lieb JD. What are super-enhancers? Nat Genet. 2015;47(1):8–12.

62. Shen Y, Yue F, McCleary DF, Ye Z, Edsall L, Kuan S, et al. A map of the cis-regulatory sequences in the mouse genome. Nature. 2012;488(7409):116–20.

63. Hay D, Hughes JR, Babbs C, Davies JOJ, Graham BJ, Hanssen L, et al. Genetic dissection of the alpha-globin super-enhancer in vivo. Nat Genet. 2016;48(8):895–903.

64. Hnisz D, Shrinivas K, Young RA, Chakraborty AK, Sharp PA. A Phase Separation Model for Transcriptional Control. Cell. 2017; 169(1): 13–23.

65. Huang J, Li K, Cai W, Liu X, Zhang Y, Orkin SH, et al. Dissecting super-enhancer hierarchy based on chromatin interactions. Nat Commun. 2018;9(1):943.

66. Gurumurthy A, Shen Y, Gunn EM, Bungert J. Phase Separation and Transcription Regulation: Are Super-Enhancers and Locus Control Regions Primary Sites of Transcription Complex Assembly? Bioessays. 2019;41(1):e1800164.

67. Bender MA, Roach JN, Halow J, Close J, Alami R, Bouhassira EE, et al. Targeted deletion of 5’HS1 and 5’HS4 of the β-globin locus control region reveals additive activity of the DNaseI hypersensitive sites. Blood. 2001;98(7):2022–7.

68. Fang X, Sun J, Xiang P, Yu M, Navas PA, Peterson KR, et al. Synergistic and Additive Properties of the Beta-Globin Locus Control Region (LCR) Revealed by 5’HS3 Deletion Mutations: Implication for LCR Chromatin Architecture. Molecular and Cellular Biology. 2005;25(16):7033–41.

69. Dukler N, Gulko B, Huang YF, Siepel A. Is a super-enhancer greater than the sum of its parts? Nat Genet. 2016;49(1):2–3.

70. Allahyar A, Vermeulen C, Bouwman BAM, Krijger PHL, Verstegen M, Geeven G, et al. Enhancer hubs and loop collisions identified from single-allele topologies. Nat Genet. 2018;50(8): 1151–60.

71. Liu J, Magri L, Zhang F, Marsh NO, Albrecht S, Huynh JL, et al. Chromatin landscape defined by repressive histone methylation during oligodendrocyte differentiation. The Journal of neuroscience: the official journal of the Society for Neuroscience. 2015;35(1):352–65.

72. Zabidi MA, Arnold CD, Schernhuber K, Pagani M, Rath M, Frank O, et al. Enhancer-core-promoter specificity separates developmental and housekeeping gene regulation. Nature. 2015;518(7540):556–9.

73. Zabidi MA, Stark A. Regulatory Enhancer-Core-Promoter Communication via Transcription Factors and Cofactors. Trends Genet. 2016;32(12):801–14.

74. Inukai S, Kock KH, Bulyk ML. Transcription factor-DNA binding: beyond binding site motifs. Curr Opin Genet Dev. 2017;43:110–9.

75. Ma KH, Svaren J. Epigenetic Control of Schwann Cells. Neuroscientist. 2018;24(6):627–38.

76. Deng W, Lee J, Wang H, Miller J, Reik A, Gregory PD, et al. Controlling long-range genomic interactions at a native locus by targeted tethering of a looping factor. Cell. 2012;149(6):1233–44.

77. Levine M, Cattoglio C, Tjian R. Looping back to leap forward: transcription enters a new era. Cell. 2014; 157(1):13–25.

78. Vietri Rudan M, Hadjur S. Genetic Tailors: CTCF and Cohesin Shape the Genome During Evolution. Trends Genet. 2015;31(11):651–60.

79. Weintraub AS, Li CH, Zamudio AV, Sigova AA, Hannett NM, Day DS, et al. YY1 Is a Structural Regulator of Enhancer-Promoter Loops. Cell. 2017; 171(7): 1573–88 e28.

80. Khan A, Fornes O, Stigliani A, Gheorghe M, Castro-Mondragon JA, van der Lee R, et al. JASPAR 2018: update of the open-access database of transcription factor binding profiles and its web framework. Nucleic acids research. 2018;46(D1):D260–D6.

81. He Y, Dupree J, Wang J, Sandoval J, Li J, Liu H, et al. The transcription factor Yin Yang 1 is essential for oligodendrocyte progenitor differentiation. Neuron. 2007;55(2):217–30.

82. He Y, Kim JY, Dupree J, Tewari A, Melendez-Vasquez C, Svaren J, et al. Yy1 as a molecular link between neuregulin and transcriptional modulation of peripheral myelination. Nature neuroscience. 2010;13(12):1472–80.

83. Guo L, Eviatar-Ribak T, Miskimins R. Sp1 phosphorylation is involved in myelin basic protein gene transcription. J Neurosci Res. 2010;88(15):3233–42.

84. Tretiakova A, Steplewski A, Johnson EM, Khalili K, Amini S. Regulation of myelin basic protein gene transcription by Sp1 and Purα: Evidence for association of Sp1 and Purα in brain. Journal of Cellular Physiology. 1999;181(1):160–8.

85. Pan MR, Hung WC. Nonsteroidal anti-inflammatory drugs inhibit matrix metalloproteinase-2 via suppression of the ERK/Sp1-mediated transcription. The Journal of biological chemistry. 2002;277(36):32775–80.

86. Bechler ME, Byrne L, Ffrench-Constant C. CNS Myelin Sheath Lengths Are an Intrinsic Property of Oligodendrocytes. Curr Biol. 2015;25(18):2411–6.

87. Snaidero N, Simons M. Myelination at a glance. J Cell Sci. 2014;127(Pt 14):2999–3004.

88. Fields RD. A new mechanism of nervous system plasticity: activity-dependent myelination. Nat Rev Neurosci. 2015;16(12):756–67.

89. Hsu PD, Scott DA, Weinstein JA, Ran FA, Konermann S, Agarwala V, et al. DNA targeting specificity of RNA-guided Cas9 nucleases. Nat Biotechnol. 2013;31(9):827–32.

90. Hwang WY, Fu Y, Reyon D, Maeder ML, Tsai SQ, Sander JD, et al. Efficient genome editing in zebrafish using a CRISPR-Cas system. Nat Biotechnol. 2013;31(3):227–9.

91. Bassett AR, Tibbit C, Ponting CP, Liu JL. Highly efficient targeted mutagenesis of Drosophila with the CRISPR/Cas9 system. Cell Rep. 2013;4(1):220–8.

92. Tuason MC, Rastikerdar A, Kuhlmann T, Goujet-Zalc C, Zalc B, Dib S, et al. Separate proteolipid protein/DM20 enhancers serve different lineages and stages of development. The Journal of neuroscience: the official journal of the Society for Neuroscience. 2008;28(27):6895–903.

93. Wang W, Kutny PM, Byers SL, Longstaff CJ, DaCosta MJ, Pang C, et al. Delivery of Cas9 Protein into Mouse Zygotes through a Series of Electroporation Dramatically Increases the Efficiency of Model Creation. J Genet Genomics. 2016;43(5):319–27.

94. Wang W, Zhang Y, Wang H. Generating Mouse Models Using Zygote Electroporation of Nucleases (ZEN) Technology with High Efficiency and Throughput. Methods Mol Biol. 2017;1605:219–30.

